# Multi-Channel Funnel Adapted Sensing Tube (MFAST) for the Simple and Duplex Detection of Parasites

**DOI:** 10.1101/2024.10.19.619249

**Authors:** Ruonan Peng, Fnu Yuqing, Taralyn J. Wiggins, Negin Bahadori, Stephen J. Dollery, Jacob Waitkus, Joshua S. Rogers, Yun Chen, Gregory J. Tobin, Ke Du

**Affiliations:** Department of Chemical and Environmental Engineering, University of California, Riverside, 900 University Ave, Riverside, CA 92507, USA; Department of Mechanical Engineering, University of California, Riverside, 900 University Ave, Riverside, CA 92507, USA; Biological Mimetics, Inc., 124 Byte Drive, Frederick, MD 21702, USA; Mechanical Engineering, College of Engineering and Science, Louisiana Tech University, Ruston, LA 71272, USA; Charlie Dunlop School of Biological Sciences, University in Irvine, California, 402 Physical Sciences Quad, Irvine, CA 92697, USA; Department of Mechanical Engineering, California State Polytechnic University, Pomona, CA, 91768, USA

## Abstract

Leishmaniasis poses a significant global health threat, infecting millions of people annually, particularly in tropical and subtropical regions. Timely and accurate detection of *Leishmania* species is crucial for effective treatment and control of this debilitating disease. This study introduces the Multi-channel Funnel Adapted Sensing Tube (MFAST) chip, an innovative diagnostic tool designed for the rapid detection of *Leishmania panamensis*. MFAST is fabricated through 3D printing and sacrificial molding of acrylonitrile butadiene styrene (ABS) and the reagents are transported between the reservoirs by gravity. We combine experiments and finite element analysis to reduce the leaking issues and facilitate smoother fluid flow, improving the overall performance of the device. Highly sensitive and specific RPA-CRISPR/Cas12a assay is utilized in the chip, achieving a detection limit as low as 1,000 parasites/mL (detecting as few as 5 parasites per reaction). The multi-channel design enables duplex detection, allowing for simultaneous identification of both *L. braziliensis* and *L. panamensis* through distinct channels. Furthermore, stability tests indicate that lyophilized reagents retain functionality for up to 15 days when stored at 4 °C, underscoring the potential of this chip for practical diagnostic applications in low-resource settings.

## Introduction

Leishmaniasis ranks second only to malaria among parasitic diseases in terms of mortality^1^. It is caused by the Leishmania parasite, a protozoan transmitted to humans through the bite of infected phlebotomine sandflies. The transmission occurs when a sandfly bites an infected person or animal and then bites another individual, injecting them with the Leishmania parasite^2,3^. The disease disproportionately affects vulnerable populations, particularly in areas with malnutrition^4^, poor housing^5^, weakened immune systems^6^, and limited healthcare access^7^. There are three main forms of leishmaniasis: cutaneous leishmaniasis (CL), visceral leishmaniasis (VL), also known as kala-azar, and mucocutaneous leishmaniasis (ML)^8^. Among these, ML can cause severe ulceration in the mucous membranes of the nose, mouth, or throat^9^. Mucocutaneous leishmaniasis rarely heals on its own and, if left untreated, can lead to life-threatening complications.

Significant facial disfigurement can occur due to the destruction of the oro/nasopharyngeal mucosa and cartilage in later stages. In severe cases, it can affect the larynx, leading to aspiration pneumonia^10^. Early detection is crucial, especially in resource-limited settings, to enable timely medical intervention and reduce the risk of disease spread^11^. While VL is the most dangerous form of the disease, it can be diagnosed using a combination of clinical signs and parasitological or serological tests^12^. However, serological tests have limited value in diagnosing CL and ML due to poor antibody responses in these cases^13^. For CL and ML, diagnosis largely relies on clinical symptoms and parasitological tests, which confirm the presence of the parasite in affected tissues^14^. These tests, however, often face challenges due to limited specificity or sensitivity^15^. Alternative nucleic acid-based approaches, such as polymerase chain reaction (PCR), offer high sensitivity but require advanced equipment and precise thermal cycling, making them less viable for point-of-care (POC) diagnostics, especially in resource-limited settings^16^.

In recent years, clustered regularly interspaced short palindromic repeats (CRISPR)-Cas 12a system has gained recognition as a promising tool for *in vitro* diagnostics, combining precision and simplicity in the detection of specific DNA sequences^17^. This system employs a guide RNA (crRNA) to recognize and cleave target DNA, a process known as *cis-cleavage*. Upon activation, it also engages in *trans-cleavage*, where it indiscriminately cuts nearby single-stranded DNA (ssDNA)^18^. This property is utilized in diagnostic assays by using ssDNA reporters tagged with a fluorophore and quencher. When Cas12a cleaves the reporter, fluorescence is generated, indicating the presence of the target DNA^19^. Unlike traditional PCR, which relies on precise thermal cycling, Cas12a operates at a constant temperature of 37 °C, making it ideal for POC applications where rapid, cost-effective, and portable diagnostics are crucial^20, 21^.

To enhance sensitivity, CRISPR-Cas12a is often coupled with isothermal amplification methods such as recombinase polymerase amplification (RPA), which can be performed at the same temperature^22^. RPA enables the rapid amplification of target DNA through the concerted actions of recombinases, single-strand binding (SSB) proteins, and DNA polymerases. The process begins when recombinases guide primers to the target DNA sequence. SSB proteins stabilize the unwound DNA strands, while DNA polymerases extend the primers along the template DNA in the presence of deoxynucleotide triphosphates (dNTPs), facilitating the synthesis of new DNA strands^23^. The RPA-CRISPR two-step process requires sequential handling to prevent premature degradation of amplified DNA, and it presents practical challenges. The requirement for manual pipetting between steps increases the risk of aerosol contamination, which can lead to false positives and complicate the workflow^24^. Our lab previously developed a funnel-adapted sensing tube (FAST) chip that combines RPA and CRISPR reactions for power-free, pipette-free, and sensitive detection of SARS-CoV-2^25^. In the FAST chip, RPA and CRISPR reagents are stored in separate chambers, isolated by carbon fiber rolls, with the reaction initiated by simply pulling the rods. This system achieved a high sensitivity of approximately 10 fM; however, the original FAST chip was a single channel and could not perform negative control on the same chip.

In this study, we introduce an all-in-one multi-channel funnel-adapted sensing tube (MFAST) chip for the duplex detection of *Leishmania* species and specific detection of *L. panamensis*, without cross-contamination. To achieve this, we perform vapor smoothing of the 3D-printed Acrylonitrile butadiene styrene (ABS) core, which increases liquid transport efficiency by approximately 10%, and the application of NeverWet coating on the valves, which reduces leakage rates by over 20%. Additionally, Plasma treatment of the inner surfaces further enhances hydrophilicity, allowing smoother fluid transport through gravity. The multi-channel design also enables simultaneous negative and positive control reactions within the same chip, reducing the likelihood of false positives and improving diagnostic reliability. The chip demonstrates the ability to detect Leishmania species with a sensitivity as low as 1,000 parasites/mL (detecting as few as 5 parasites per reaction), enabled by a highly specific RPA-CRISPR/Cas12a assay. By integrating a portable blue light transilluminator, this chip offers a power-free, pipette-free solution, significantly simplifying on-site diagnostics. The duplex testing capability further enhances efficiency by minimizing reagent consumption and enabling the simultaneous detection of multiple species. This versatile, low-cost platform is well-suited for POC diagnosis of parasites, particularly in low-resource settings.

## Results and Discussion

The fabrication process of the MFAST chip is illustrated in **Fig. 1a**. The chip core was 3D-printed using ABS material. To create a smooth surface, the printed core was suspended in a vapor chamber with heated acetone, which dissolved surface imperfections and produced a glossy finish. The smoothed core was then attached to a petri dish, and a mixture of polydimethylsiloxane (PDMS) and curing agent was poured into the mold. Once cured, the chip was submerged in acetone, dissolving the ABS core and leaving behind the final microfluidic structure. **Fig. 1b** shows a photograph of a fabricated MFAST chip filled with food dye, which is distinctly observable to the naked eye. The chambers are delineated by 3D-printed rectangle pieces that serve as functional valves. The volumes of the chambers, from top to bottom, are around 338 μL, 160 μL, and 160 μL, respectively. The corners of the chambers are rounded to minimize dead volume, thereby enhancing the efficiency of reagent flow. This design modification facilitates smoother transport of reagents, ultimately improving the overall operational efficiency of the chip.

**Fig. 1.**
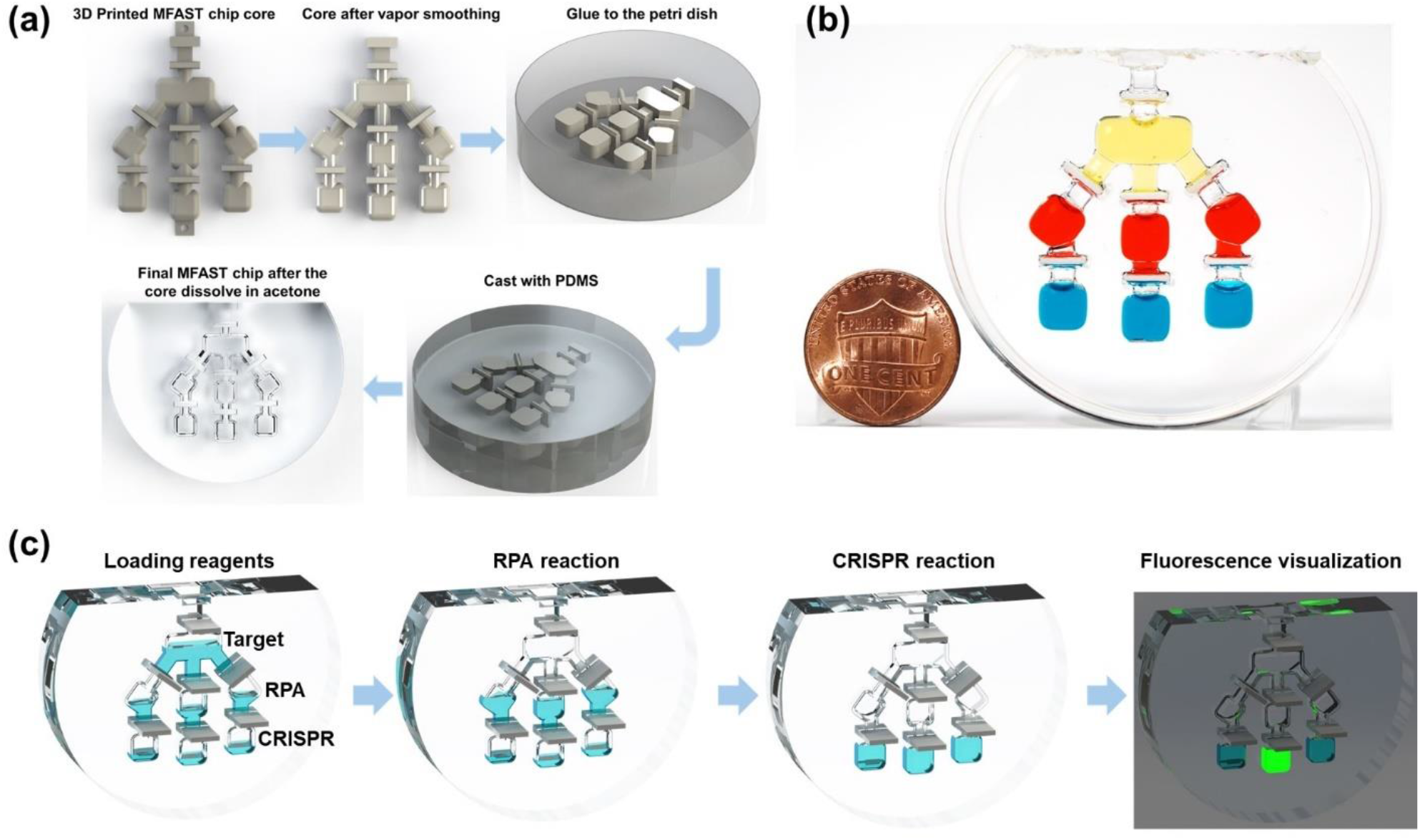
Schematic of MFAST chip. (a) Fabrication process of MFAST chip. (b) Photograph of MFAST filled with food dye with a US one-cent coin for scale. (c) Working principle of the MFAST for visual detection of *Leishmania*.

**Fig. 1c** illustrates the visual detection process employed by the self-contained MFAST chip. Prior to initiating the reaction, RPA and CRISPR reagents were systematically loaded into the designated chambers of the MFAST chip. Specifically, 26.5 μL of the CRISPR-Cas12a complex was introduced into the bottom chambers, while a total of 68.6 μL of a mixture containing primers and a rehydrated TwistAmp^@^ basic reaction pellet was loaded into the second row of the chamber. The top chamber was filled with 75 μL of a nuclease-free water and magnesium acetate mixture, which is used to activate the RPA reaction. Once 15 μL of the target was added to the top chamber, the chip was prepared for visual detection. The amplification reaction commenced by pulling the top valves, enabling the target to flow downwards and mix with the RPA reagents. After a 20-min incubation at 37 °C on a heating pad, the resulting amplicons were transferred to the CRISPR reaction chamber for an additional incubation of 30 min at the same temperature. Following this, a blue light transilluminator was utilized to excite the fluorophore, allowing for naked-eye observation of the reaction results. This methodological approach underscores the integration of RPA and CRISPR technologies within the MFAST chip framework, highlighting its potential for efficient and accessible visual detection in various applications.

After fabricating the MFAST chip, surface treatments were made to optimize its performance, including the application of a NeverWet coating on the valves and vapor smoothing of the 3D-printed core. To demonstrate the leak-prevention capabilities of the NeverWet coating, **Fig. 2a** shows a food dye test. In the left image, the MFAST chip is loaded with yellow, red, and blue dyes in the top, middle, and bottom chambers, respectively. The valves in the left and middle channels were coated with NeverWet, while the right channel remained uncoated. In the middle image, after opening the second row of side valves and shaking the chip, the yellow and red dyes were mixed to form an orange in the coated left channel. In contrast, in the uncoated right channel, the yellow dye leaked into the red, followed by mixing with the green dye. In the right image, after pulling the bottom side valves, the orange dye mixed with the green in the coated left channel. Meanwhile, in the uncoated right channel, all the dyes were mixed. The valves in the middle channel, sealed with the NeverWet coating, prevented any leakage or mixing.

**Fig. 2.**
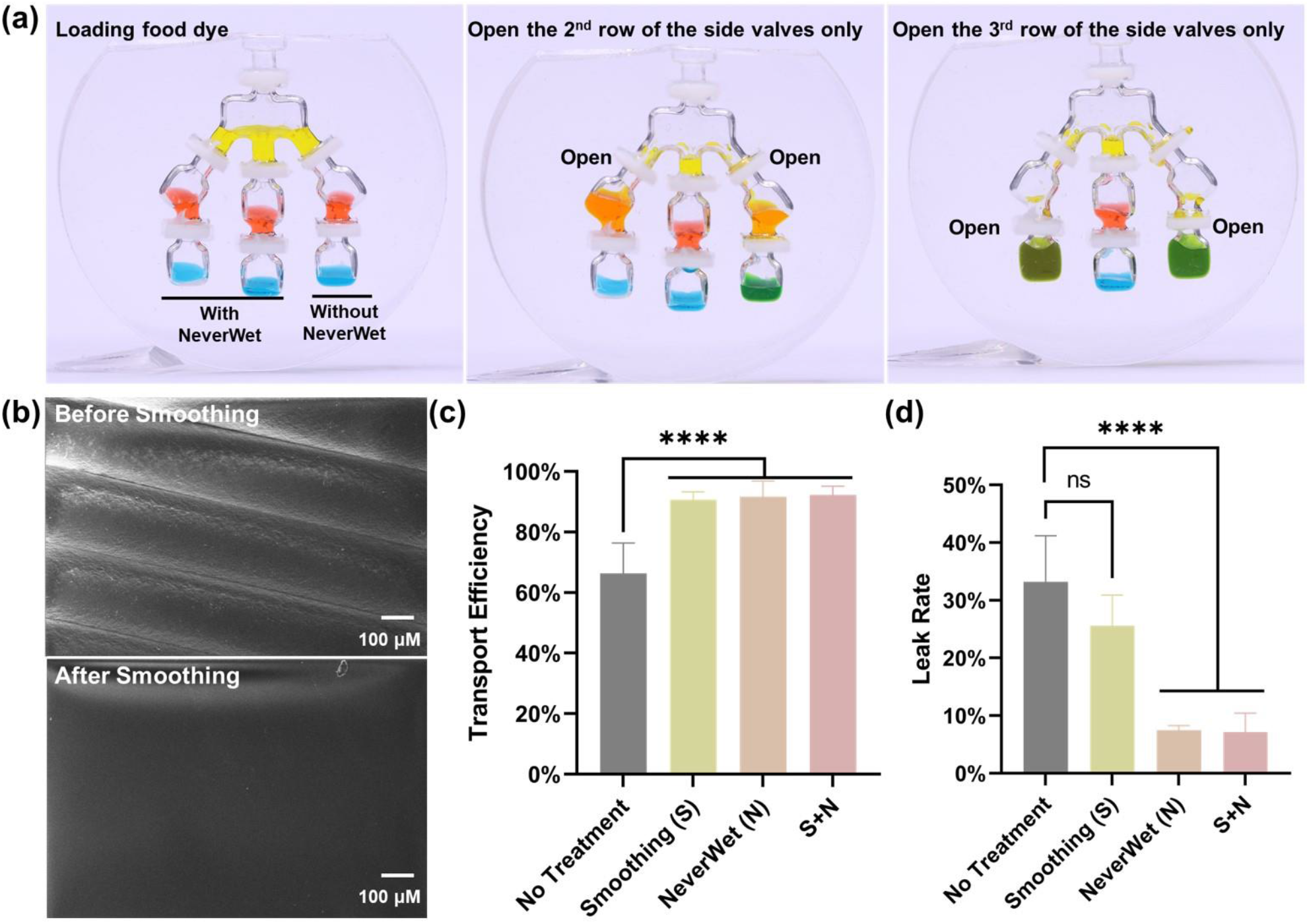
Surface treatment and quantification of MFAST chip. (a) Food dye test evaluating the leak-prevention capabilities of the NeverWet coating. The left photo illustrates the food dye loading process with all valves closed. The valves in the left and middle channels were coated with NeverWet, while the right channel remained uncoated. The middle photo depicts the mixing of food dye after opening only the second row of side valves while keeping the valves of the middle channel closed. The right photo shows the mixing of food dye after opening only the valves on the bottom row while keeping the valves of the middle channel closed. (b) SEM images of the PDMS replica of the ABS core before (top) and after (bottom) vapor smoothing. (c) Transport efficiency of the MFAST chip without treatment, with vapor smoothing of the ABS core, with NeverWet coating on the valves, and with both treatments combined. (d) The leak rate of the MFAST chip without treatment, with vapor smoothing of the ABS core, with NeverWet coating on the valves, and with both treatments combined. Statistical analyses were performed using one-way ANOVA analysis. The data are represented as mean ± standard deviation (*n = 3*). ns, not significant = *p>0.05; * =0.01< p*≤*0.05; ** =0.001< p*≤*0.01; **** = p*≤*0.0001*.

The vapor smoothing would eliminate the layer lines created during the 3D printing process and result in a glossy finish on the core surface. This process transformed the replicated PDMS surface from translucent to transparent, which improved visibility for users to observe the status of the chambers. As shown in **Fig. 2b**, evaluated under SEM, vapor smoothing not only improved the visual appearance but also enhanced the performance of the MFAST chip. **Fig. 2c** shows that this process increased the transport efficiency of liquids by reducing the layer lines that could trap fluid during transport. The efficiency of liquid transport was tested by measuring the amount of food dye transferred from the second row of the MFAST chip to the third row. Results demonstrated that vapor smoothing improved transport efficiency by approximately 10%. The NeverWet coating, by creating a superhydrophobic surface, reduced liquid adhesion to the valve and repelled fluid trapped in the intricate design of the valves. NeverWet played a crucial role in the leak prevention test of the MFAST chip. In this test, food dye was added to the second row of the chip, and the amount of remaining dye was measured after shaking the chip five times with all valves closed. The results, shown in **Fig. 2d**, revealed that the NeverWet coating reduced leakage by more than 20%, providing significant protection against leaks. While vapor smoothing had a less substantial effect on leak prevention, it still provided slight improvements in the absence of NeverWet.

To improve liquid transfer, the water contact angle of the inner MFAST chip chamber was altered. Our simulation, as shown in **Fig. 3a**, illustrates that lowering the water contact angle to 30°, results in the easiest flow pattern. With that in mind, oxygen plasma treatment was used to decrease the water contact angle of the MFAST chip’s inner surface^26^. As shown in **Fig. 3b**, the water contact angle of the inner cavity surface dropped from 106.05° to 39.09° after the plasma treatment. We found that the transportation of the fluid to the subsequent chamber became much easier after the plasma treatment with only a little tap needed.

**Fig. 3.**
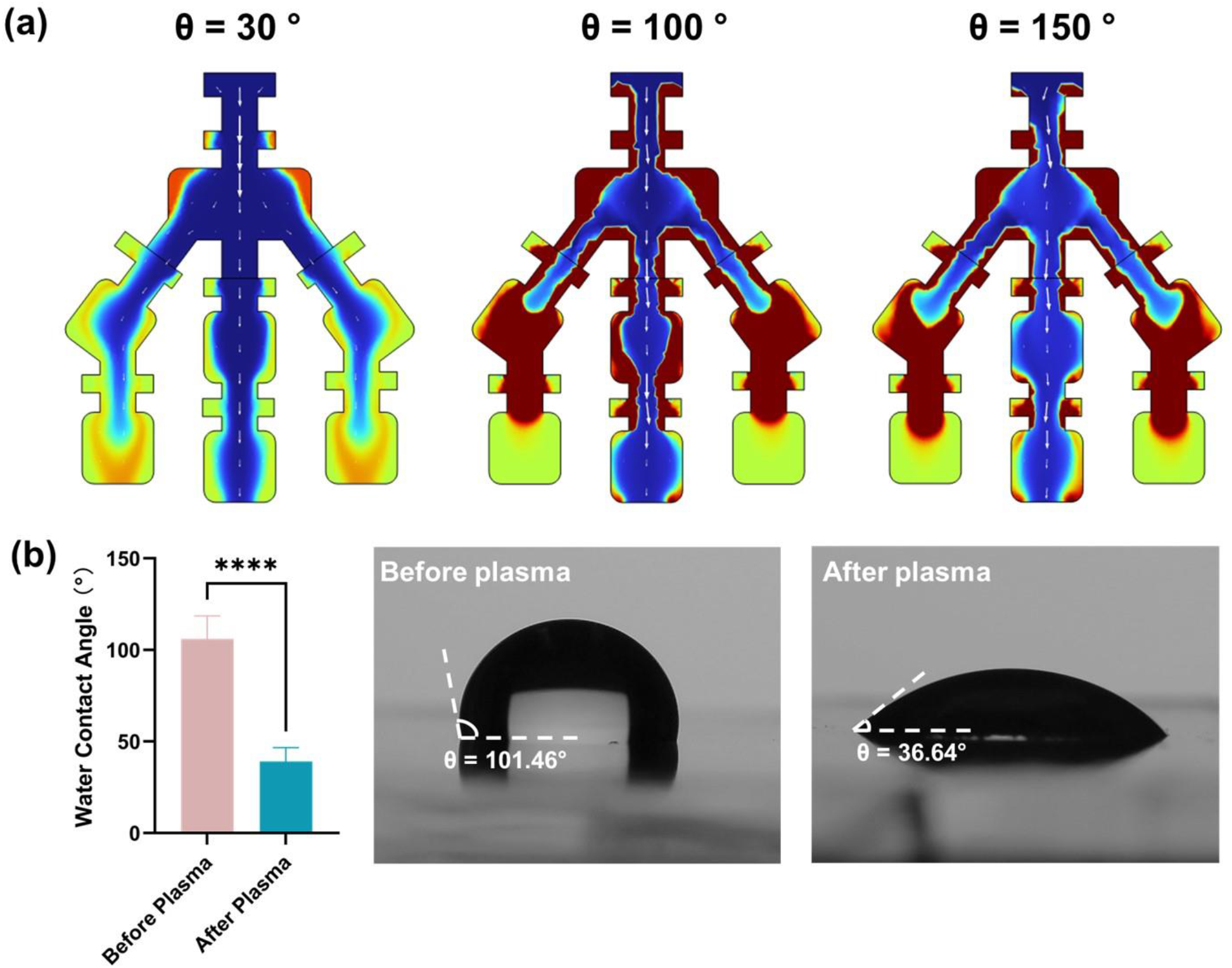
Liquid flow simulation and effect of plasma treatment on wettability. (a) Simulation results of volume fraction showing the effect of varying surface wettability at time = 6 s. (b) Static contact angles of MFAST chip before (left) and after (right) plasma treatment. Statistical analyses were performed using unpaired t-test analysis. The data are represented as mean ± standard deviation (*n = 3*). ns, not significant = *p > 0.05; * =0.01 < p*≤*0.05; ** =0.001< p*≤*0.01; **** = p*≤*0.0001*.

The specificity and sensitivity of the RPA-CRISPR assay for the detection of *L. panamensis* were subsequently evaluated. To assess specificity, fluorescence was compared between *L. panamensis* at a concentration of 1 × 10^6^ parasites/mL and six other *Leishmania* strains, namely *L. braziliensis, L. gerbelli, L. infantum, L. tropica, L. donovani*, and *L. venezuelensis*, as well as a non-template control (NTC). The endpoint fluorescence images of the reactions are shown in **Fig. 4a**. Upon excitation with a blue light transilluminator, the fluorescence differences between *L. pana-mensis* and the other groups are clearly distinguishable by the naked eye. The normalized fluorescence signal is presented in **Fig. 4b**, indicating that the fluorescence intensity of the six non-target strains and the NTC group was significantly lower than that of *L. panamensis*, demonstrating the high specificity of the RPA-CRISPR assay. To determine the detection limit, different concentrations of *L. panamensis* were used as targets and added to the RPA mixture. Following the RPA reaction, the amplicons were introduced into a pre-mixed CRISPR reaction and incubated at 37 °C for 30 min. For the negative control groups, either the RPA primers or the crRNAs were replaced with nuclease-free water. The endpoint fluorescence images of the reaction are shown in **Fig. 4c**, with the corresponding normalized fluorescence signals presented in **Fig. 4d**. Distinct fluorescence differences were observed at concentrations of 1 × 10^4^ and 1 × 10^3^ parasites/mL, whereas no significant fluorescence difference between positive and negative groups was detected at a concentration of 1 × 10^2^ parasites/mL or lower. The on-chip detection results are presented in **Fig. 4e**. The left channel represents the negative group without RPA primers, while the right channel represents the positive group without crRNAs. Compared to the negative groups, the positive group exhibited a significantly stronger fluorescence signal at a concentration of 1 × 10^3^ parasites/mL, demonstrating consistency with the off-chip results.

**Fig. 4.**
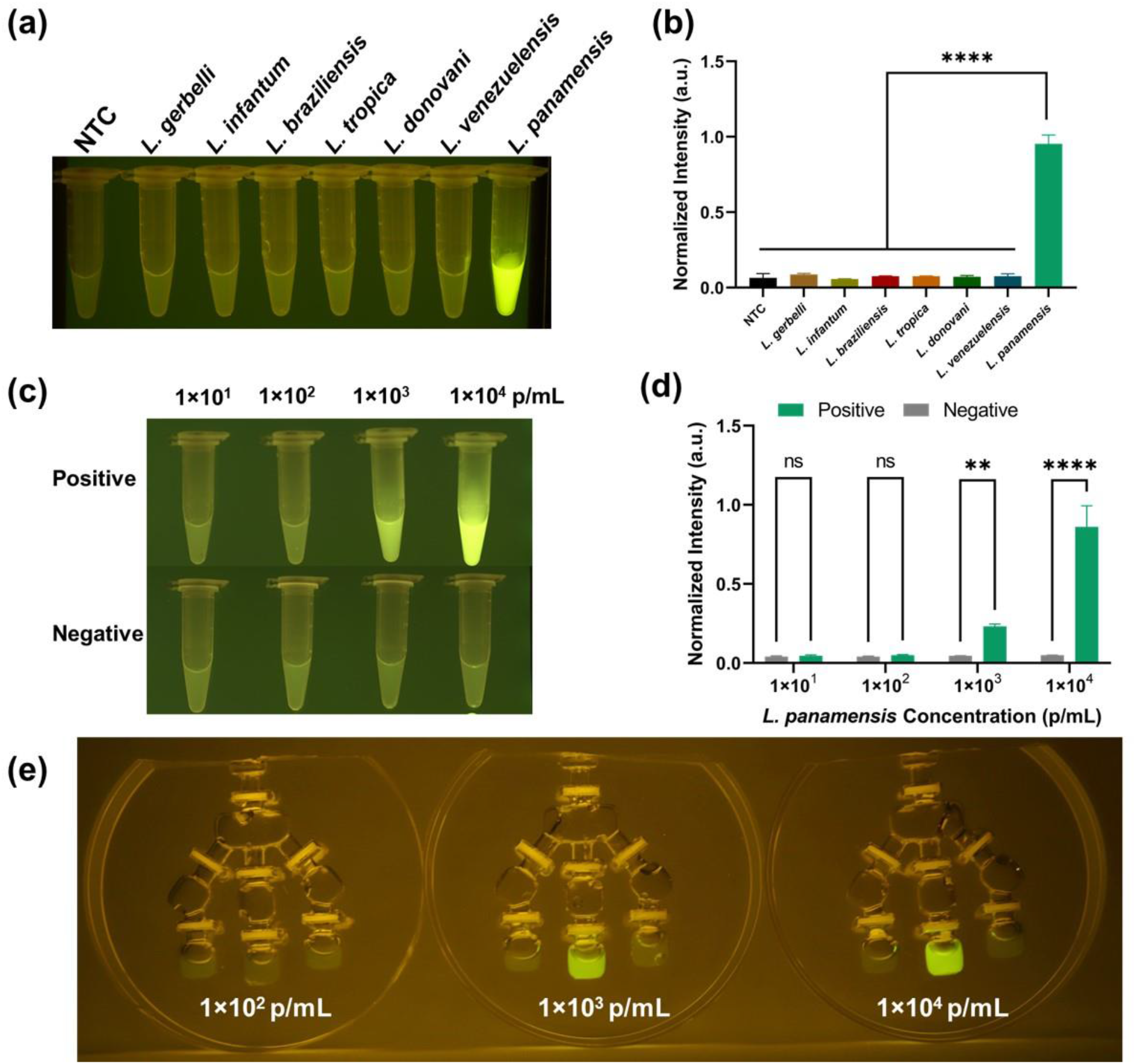
(a) Endpoint images of the reactions with 1 × 10^6^ parasites/mL of different *Leishmania* strains. Fluorescence was excited by a transilluminator (wavelength: 465 nm) (b) Normalized fluorescence signals of the reactions with 1 × 10^6^ parasites/mL of different *Leishmania* strains. Statistical analyses were performed using one-way ANOVA analysis. (c) Endpoint images of the reactions excited by a transilluminator (excitation wavelength: 465 nm) in response to different *L. panamensis* concentrations ranging from 10 to 1 × 10^4^ parasites/mL. (d) Normalized fluorescence signals of the reactions with different *L. panamensis* concentrations ranging from 10 to 1 × 10^4^ parasites/mL. Statistical analyses were performed using two-way ANOVA analysis. (e) Endpoint images of the on-chip reactions excited by a transilluminator (excitation wavelength: 465 nm) in response to different *L. panamensis* concentrations ranging from 100 to 1 × 10^4^ parasites/mL. “NTC” refers to no template control. The data are represented as mean ± standard deviation (*n = 3*). ns, not significant = *p > 0.05; * =0.01 < p*≤*0.05; ** =0.001< p*≤*0.01; **** = p*≤*0.0001*.

Since the chip was designed with multiple channels, it can be applied for duplex detection by loading two sets of RPA primers and crRNAs targeting different *Leishmania* species. One channel was used for the detection of all seven *Leishmania* species, while another channel was specific for *L. panamensis*, which is associated with mucocutaneous diseases. The third channel served as a negative control without RPA primers. Among the seven species, *L. braziliensis* was used as the sole target, with the CRISPR and RPA reagents for detecting all seven species loaded into the right channel, while the left two channels served as negative controls. The results presented in **Fig. 5a** (left) indicate that the positive group generated noticeably stronger fluorescence compared to the negative controls, demonstrating effective detection of the target without cross-contamination and false-positive results. When a 1:1 mixture of *L. panamensis* and *L. braziliensis* was analyzed, with the RPA and CRISPR reagents specific to *L. panamensis* loaded into the left channel of the MFAST chip. As shown in **Fig. 5a** (right), both the left and right channels displayed strong green fluorescence, whereas the middle negative group did not, thereby confirming the successful duplex detection of both species. The quantitative normalized fluorescence intensity in **Fig. 5b** further supports these findings, showing that the *Leishmania* groups exhibited considerably higher fluorescence intensity compared to the negative control.

**Fig. 5.**
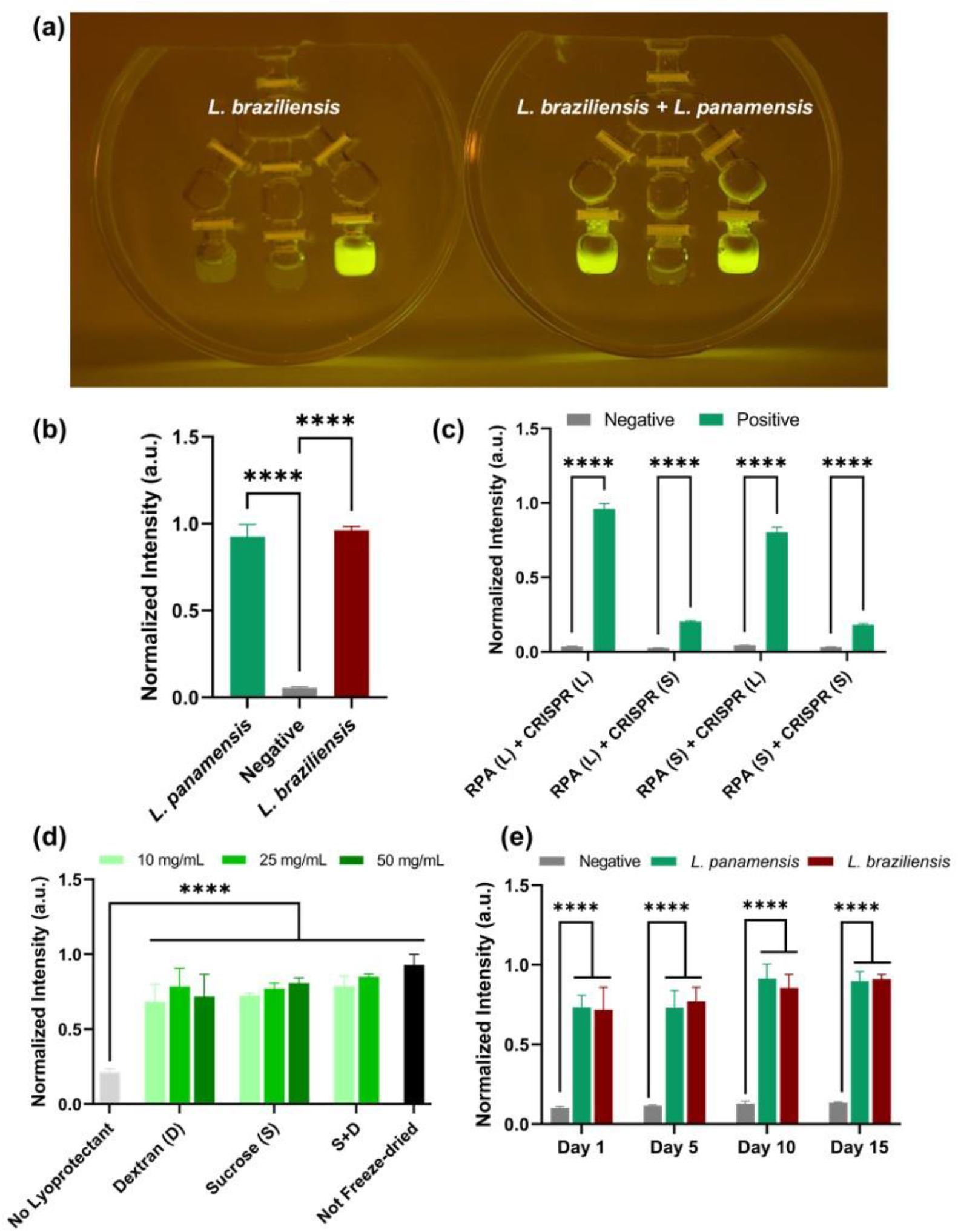
Duplex detection on MFAST. (a) Endpoint images of the on-chip reaction with 1 × 106 parasites/mL of *L. braziliensis* (left) and a 1:1 mixture of *L. panamensis* and *L. braziliensis*. In the left image, the two left channels represent negative controls. In the right image, the middle channel serves as a negative control. Fluorescence was excited by a transilluminator (wavelength: 465 nm) (b) Normalized fluorescence signals from the reactions with 1 × 10^6^ parasites/mL of a 1:1 mixture of *L. panamensis* and *L. braziliensis*. Statistical analyses were performed using one-way ANOVA analysis. (c) Impact of lyophilization on the performance of RPA and CRISPR reactions. (d) Performance of lyophilized reactions supplemented with dextran and/or sucrose at concentrations of 10, 25, and 50 mg/mL. (e) Stability test of lyophilized reactions stored at 4 °C. Statistical analyses for lyophilization results were performed using two-way ANOVA analysis. The data are represented as mean ± standard deviation (*n=3*). ns, not significant = *p>0.05; * =0.01< p*≤*0.05; ** =0.001< p*≤*0.01; **** = p*≤*0.0001*.

To enhance the storage stability^27^, ease of transportation^28^, and overall usability of the MFAST chip, the reagents were subjected to lyophilization. This process involves initially freezing the reagents, followed by sublimation of water during the primary drying phase. Secondary drying is then conducted to eliminate residual moisture and achieve the target final moisture content^29^. The RPA and CRISPR reagents were lyophilized independently, and their performance was subsequently evaluated and compared. The enzyme pellet in the RPA kit, which was originally supplied in lyophilized form, was added after the RPA reagents underwent the freeze-drying process. The input target was a 1:1 mixture of *L. panamensis* and *L. braziliensis* with a concentration of 1 × 10^6^ parasites/mL. The results, as shown in **Fig. 5c**, demonstrate that the fluorescence intensity generated by the lyophilized solid form (S) was significantly lower than that of the liquid reagents (L), indicating a reduction in assay performance. To protect Cas12a during freeze-drying, stabilizing agents like dextran and/or sucrose were added to maintain its structure and functionality^30^. As shown in **Fig. 5d**, we found that the activity of reactions supplemented with sucrose individually increased with concentration, but dextran exhibited a concentration optimum of 25 mg/mL in the final reaction. Combining 25 mg/mL each of sucrose and dextran exhibited approximately 85% recovery of the pre lyophilization signal. Additionally, we assessed the stability of the lyophilized reagents stored at 4 °C. As depicted in **Fig. 5e**, the lyophilized reagents remained functional and stable for up to 15 days.

The freeze-drying process negatively affected the efficiency of both RPA and CRISPR reactions, with a more pronounced reduction observed for CRISPR, likely due to the loss of Cas12a activity during lyophilization. This loss can be attributed to stresses induced during the lyophilization process, including cold denaturation, exposure to ice-water interfaces, and freeze concentration, which can lead to protein unfolding or aggregation. During the drying phases, the removal of water also affects its structural integrity, potentially causing irreversible denaturation^31^. To enable a shelf-stable MFAST system for POC applications, excipients such as dextran and sucrose are applied in our assay to form hydrogen bonds with specific sites on the protein surface, effectively replacing the thermo-dynamic stabilization role of water that is lost during drying. This process increases the free energy of protein unfolding, thereby preventing denaturation and preserving the native structure, ultimately enhancing storage stability^32^.

Compared to the previous FAST chip^25^, the chip design has changed from one column to three columns for duplex detection. The valve design was changed from a carbon fiber rod to a 3D-printed flat valve with NeverWet coating to enhance the sealing between the chambers so that the FAST chip no longer needs to be cut into two pieces. The material for the MFAST sacrificial core has changed from polyvinyl alcohol (PVA), water-soluble, to acrylonitrile butadiene styrene (ABS), acetone-soluble, due to its better printability, cheaper material cost, and faster dissolving time. Using 3D-printed ABS for PDMS molding has been reported by many studies, and it has been proven to be a promising way to produce PDMS replicas^33-37^. Our new process also improves the visibility of the chip by using the vapor smoothing process to remove the layer lines created from the 3D printing process, which has been proven to be an effective method to polish the surface of the 3D-printed parts^33^. Plasma treatment of the chamber was added to improve the fluid flow between the chambers. Overall, the vapor smoothing, NeverWet coating, and plasma treatment were to improve the usability and reliability of the MFAST chip, which is an important step toward real-world applications.

This multi-channel setup enables both positive and negative results to be observed simultaneously on a single chip by simply pipetting the target into the reagent-preloaded system, streamlining the detection process for *L. panamensis* and minimizing the risk of false positives. By integrating both positive and negative reactions into a closed system, the chip significantly reduces contamination risks, simplifies handling, and enhances its suitability for rapid POC diagnostics. Moreover, the duplex detection design allows for the simultaneous detection of different *Leishmania* species within a single run, increasing throughput and shortening assay time compared to sequential testing. This not only conserves reagents by running multiple assays on the same chip, lowering overall costs but also improves result reliability through integrated negative controls for real-time comparison, further reducing the chances of false positives.

The other detection and quantification methods of the parasitic such as microscopy^38^, Raman spectrometry^39^, and quartz crystal microbalance^40^ would require trained technicians and expensive equipment. On the other hand, the MFAST device only requires a heating device and a UV flashlight and does not require a lab environment to perform the test allowing it to be used in POC applications. Users can quickly determine the result based on the fluorescence signal without the need for a detection device or training^41,42^.

To achieve higher multiplexing capability and easier fluid transport, in the future, the inner surface can be decorated with microstructures such as superhydrophilic honeycomb structure^43^. During the testing of the chip, it was observed that the distribution of the liquid when transporting from the top row of the chamber to the middle row was not even, where the middle column would have more liquid than the other two columns. In the future, the diameter of the channel could be changed with the help of simulation to produce an even distribution of the liquid across all the chambers and allow it for more than three columns.

## Conclusions

We developed an all-in-one MFAST chip using sacrificial molding for *in vitro* detection of *L. panamensis*. Several surface treatments were applied to enhance its performance. Vapor smoothing of the ABS core improved liquid transport efficiency by 10% while coating the valves with NeverWet reduced leakage by over 20%. Additionally, plasma treatment was performed on the inner surfaces of the MFAST chip to increase hydrophilicity, facilitating smoother liquid flow, as demonstrated in simulation results. For detection, we employed the RPA-CRISPR/Cas12a assay, which showed high specificity for *L. panamensis* compared to other *Leishmania* strains, along with a sensitivity of 1,000 parasites/mL. To expand its diagnostic capabilities, the multi-channel design of MFAST enabled duplex detection by incorporating distinct primers and crRNAs in separate channels. This successfully achieved simultaneous detection of *L. braziliensis* and *L. panamensis* without cross-contamination. Furthermore, we tested the stability of lyophilized reagents preloaded in the chip. With excipients of 25 mg/mL of sucrose and dextran, the lyophilized reagents remained functional and stable for up to 15 days. The MFAST chip demonstrates a promising platform for leishmaniasis diagnosis, offering the potential for multiplexing and reagent stability in field settings.

## Experiment Methods

### Fabrication of MFAST Chip

The 3D model of the MFAST chip core was designed using SolidWorks. It was fabricated using fused deposition modeling X1 Carbon 3D printer (Bambu Lab) with PolyLite™ ABS (Polymaker) material. The printed core was then suspended in the vapor smoothing chamber with a copper wire for two and a half minutes to remove the layer lines. The vapor smoothing chamber consists of a tall glass jar with approximately 1 cm of acetone covering the bottom and was heated at 90 °C. During this period, the vapored acetone would dissolve the surface of the printed ABS core therefore removing the layer line and creating a glossy finish, as shown in Fig. 2a. The hooks, designed for hanging by the wire, were snipped off after the smoothing process. It was then glued on the 60 × 15 mm petri dish with super glue (Loctite). The 10:1 ratio mixture of the Polydimethylsiloxane (SYLGARDTM 184 Silicone Elastomer Kit) and curing agent was poured into the petri dish and filled to the top. It was then placed at room temperature for 2 days to degas before being placed into an oven at 70 °C for 4 hours. After curing, the MFAST chip was submerged with acetone in an ultrasonic cleaner until the ABS core was fully dissolved. The valves of the MFAST chip were 3D printed using UltraCraft Reflex 3D Printer combo (HeyGears) with UltraPrint-Modeling PAT10 Transparent Resin (HeyGears). Then, the surface of the valve was coated with NeverWet (Rust-Oleum, No. 274232). The valves were left to dry for 30 min in between the two coating steps. Both steps were applied from approximately 20 cm away, and then air dried at room temperature overnight before use.

### Surface treatment and quantification

The transport efficiency of the chip was measured by adding 100 μL of food dye into the chambers in the second row and then extracting the food dye from the third row after it had been transported and repeated 6 times. The weight of the food dye was measured by using an electronic balance. The leak test was performed by adding 100 μL of food dye into the second row closing all the valves then shaking it 5 times by hand. Then, the liquid remaining in the second row was extracted and measured. The water contact angle of the inner chamber was measured by creating a large flat cavity surface using the same process as the MFAST chip with ABS sacrificial molding. It was then cut into size for the water contact angle testing. The plasma treatment of the MFAST chip was performed by placing it into an oxygen plasma chamber for 10 min with 100 W RF power, 20 sccm O_2_, and 0.6 mBar pressure.

### Two-phase flow simulation

The simulation investigates a two-phase flow involving water as the continuous phase and air as the dispersed phase, using COMSOL Multiphysics. The primary objective is to model the interaction between the dispersed air phase and the continuous water phase in a controlled geometry. The Laminar Flow interface uses the Navier-Stokes equation for incompressible fluid flow, ensuring that the velocity and pressure fields of the continuous water phase are accurately simulated. The Phase Transport interface models the behavior of the dispersed air phase, starting from a closed valve configuration filled with air and transitioning to an open valve, allowing water to flow. Boundary conditions include an inlet velocity defined by a step function, allowing a gradual increase in velocity from 0 m/s to 0.01 m/s. This simulates valve opening. The initial conditions set the water as uniformly distributed, with air present at a volume fraction of 0.8, representing the initial state with closed valves. The simulation explores different contact angles (30°, 100°, and 150°) to observe changes in the volume fraction and velocity fields.

### Parasite culture and lysis

The following *Leishmania* species were purchased from BEI Resources: *L. panamensis*, strain PSC-1 (MHOM/PA/94/PSC 1), NR-50162; *L. braziliensis*, strain HOM/BR/75/M2903, NR 50608; *L. gerbelli*, strain RHO/CN/62/20, NR-50601; L *infantum*, strain HOM/CN/93/KXG-LIU, NR 50605; *L. tropica*, strain HOM/TR/99/EP41, NR-51828; *L. donovani*, strain HOM/IN/83/AG83, NR-50602; *L. venezuelensis*, strain MHOM/VE/80/H-16, NR-29184. The parasites were cultivated in tissue culture by inoculating frozen samples into T-25 flasks containing Modified M199 medium (Gibco, Ref: 12350-039) supplemented with 10% heat-inactivated fetal bovine serum (Neuromonics, Cat. No. FBS006) and 10 μg/mL hemin chloride (Millipore Sigma, Ref: 3741-5GM) at 25 °C, and subsequently expanded into T-182 flasks. Parasite density was determined using a cytometer, followed by pelleting at 2,000 × g and washing with PBS. At peak density, parasites were cryopreserved in 0.5 mL aliquots at approximately 3 × 10^7^ parasites/mL with 10% DMSO and stored in liquid nitrogen for long-term preservation. For parasite crude lysis, 12 mL of peak-density parasites were harvested, pelleted, washed with 10 mL of PBS, and aliquoted into 1.5 mL Eppendorf tubes. The tubes were centrifuged to form a pellet, and 200 μL of a 0.1% Triton X-100 solution was added, followed by incubation at 56 °C for 10 min.

### RPA amplification and CRISPR-Cas12a detection

The TwistAmp^@^ Basic kit was obtained from TwistDx™. NEBuffer™ r2.1 was purchased from New England Biolabs. The RPA primers, fluorophore-quencher (F-Q) probe, crRNA, and AsCas12a (Alt-R™ A.s. Cas12a Ultra) were purchased from Integrated DNA Technologies, with detailed information about the synthetic oligonucleotides provided in Table S1. The DNA sequence of different *Leishmania* species was aligned using clustalW, and the RPA primer sets were designed using the Primer 3 program.

The target mixture was prepared by combining 5 μL of the target with 21.25 μL nuclease-free water and 3.75 μL of magnesium acetate (280 mM). To prepare the RPA mixture, 4.8 μL of RPA primers (10 μM) were combined with 59 μL of primer free rehydration buffer, and the TwistAmp^@^ Basic reaction pellet was dissolved in the mixture. For the CRISPR mixture, 7.5 μL of crRNA (1 μM) was mixed with 6 μL of Cas12a (1 μM), and the complex was then equilibrated at room temperature for 10 min. Subsequently, 12 μL of NEBuffer™ r2.1 (10×) and 1 μL of ssDNA probe (100 μM) were added to complete the CRISPR mixture.

The target mixture was added to the RPA mixture, where the magnesium acetate contained in the target mixture initiated the RPA reaction. The reaction was incubated at 37 °C for 20 min. Following this, the mixture was added to the pre-assembled CRISPR mixture and incubated at 37 °C for 30 min.

### Leishmania detection with MFAST

The CRISPR mixture was pipetted into the bottom chamber, and valves were inserted into bottom gaps to seal the CRISPR reagents. Subsequently, the RPA mixture was introduced to the second chamber, and the valves were inserted into the corresponding gaps to seal the RPA reagents. The target mixture was then introduced to the top chamber and sealed with valves. For negative controls, either the RPA primers or the crRNAs were replaced with nuclease-free water. To initiate the reaction, the second row of valves was pulled to allow the target mixture to flow into the RPA chamber. The MFAST chip was then incubated on a heating pad at 37 °C for 20 min. Following this incubation, the bottom valves were pulled to allow the mixture to flow into the CRISPR chamber, which was incubated at 37 °C for an additional 30 min. A blue light transilluminator (SmartBlue, Part No: NEB-E4100, excitation wavelength of 465 nm) was subsequently used to excite the mixture for naked-eye observation. Finally, 20 μL of nuclease-free water was added to 5 μL of the mixture, and the resulting solution was characterized using an Agilent BioTek Cytation 5 imaging reader.

### Lyophilization

For the off-chip assay, given that the RPA enzymatic pellets were provided in a lyophilized form, the lyoprotectant dextran and/or sucrose was incorporated into the CRISPR reaction prior to transferring the mixture to PCR strip tubes, each with a small hole punctured in the cap. When using a 384-well plate, a PCR sealing film (Thermo Fisher Scientific) was applied over the wells, with a small hole punctured in each well. The RPA and CRISPR reactions were subsequently pre-frozen and freeze-dried in a freeze dryer (Harvest Right, Model No: HRFD-SML-BK) overnight. For the on-chip lyophilization, following the loading of reagents into the respective chambers, the chip was laid flat with the valves slightly open to leave a gap, and then pre-frozen and freeze-dried in a freeze dryer overnight.

## Author Contributions

Ruonan Peng, FNU Yuqing, Gregory J. Tobin and Ke Du designed the experiments. Ruonan Peng, FNU Yuqing, Taralyn J. Wiggins, Negin Bahadori, Stephen J. Dollery, Jacob Waitkus, Joshua S. Rogers performed the experiments. Ruonan Peng and FNU Yuqing wrote the manuscript. All authors commented on the manuscript.

## Acknowledgements

This project was supported by NIH NIAID and University of California OASIS Internal Grant.

## Data Availability

The data supporting the results in this study are available within the paper. Further information can be requested from the corresponding authors via e-mail.

